# Trapping virus-loaded aerosols using granular protein nanofibrils and iron oxyhydroxides nanoparticles

**DOI:** 10.1101/2022.06.29.498082

**Authors:** Antonius Armanious, Heyun Wang, Peter A. Alpert, Chiara Medaglia, Mohammad Peydayesh, Arnaud Charles-Antoine Zwygart, Christian Gübeli, Stephan Handschin, Sreenath Bolisetty, Markus Ammann, Caroline Tapparel, Francesco Stellacci, Raffaele Mezzenga

## Abstract

The ongoing COVID-19 pandemic has revealed that developing effective therapeutics against viruses might be outpaced by emerging variants,^1–5^ waning immunity,^6–9^ vaccine skepticism/hesitancy,^10–12^ lack of resources,^13–16^ and the time needed to develop virus-specific therapeutics,^17,18^ emphasizing the importance of non-pharmaceutical interventions as the first line of defense against virus outbreaks and pandemics.^19–23^ However, fighting the spread of airborne viruses has proven extremely challenging,^23–28^ much more if this needs to be achieved on a global scale and in an environmentally-friendly manner.^29,30^ Here, we introduce an aerosol filter made of granular material based on whey protein nanofibrils and iron oxyhydroxides nanoparticles. The material is environmentally-friendly, biodegradable, and composed mainly of a dairy industry byproduct.^31^ It features remarkable filtration efficiencies between 95.91% and 99.99% for both enveloped and non-enveloped viruses, including SARS-CoV-2, the influenza A virus strain H1N1, enterovirus 71, bacteriophage Φ6, and bacteriophage MS2. The developed material is safe to handle and recycle, with a simple baking step sufficient to inactivate trapped viruses. The high filtration efficiency, virtually-zero environmental impact, and low cost of the material illuminate a viable role in fighting current and future pandemics on a global scale.

---

The development of vaccines and other therapeutics is an essential component of our panoply to fight viral pandemics. Therapeutics alone, however, might not be sufficient to end a pandemic with the desired speed, leading to potentially avoidable fatalities^32,33^ and adverse socioeconomic consequences.^34^ The emergence of new variants that can evade the immune response of vaccinated and convalescent patients,^1–5^ waning immunity,^6–9^ vaccine skepticism/hesitancy,^10–12^ and lack of resources for production and administration of vaccines on a global scale within a short time period^13–18^ are all factors that might compromise efforts to end a pandemic through therapeutics. Therefore, a key tool in this fight is preventing the transmission of viruses through non-pharmaceutical interventions.^19–23^ It has long been thought that the transmission of airborne viruses is mainly driven by droplets;^25,35^ a growing body of evidence reveals that aerosols substantially contribute to airborne viral transmission, particularly in indoor spaces.^25,35–38^ Mask mandates, social distancing, increased ventilation, and the use of air filters are measures used to combat airborne virus transmission in indoor spaces. Introducing air filters offers several advantages over other measures, i.e., they could contribute to maintaining indoor space capacities, offer cost-effective solution for poorly ventilated spaces, and are less sensitive to personal choices and/or behavioral discipline. High-efficiency particulate air (HEPA) filters constitute the gold standard for the filtration of aerosols. Producing HEPA filters requires relatively advanced fabrication technologies,^39^ with a large proportion made of either glass fibers or plastics, the former of which requires an energy-intensive fabrication process, and the latter of which relies heavily on the petrochemical industries.^39,40^ In addition, over time, a filter cake builds up on the fibers of the filter, resulting in increased resistance to airflow and the imminent need to replace the whole filter, with very limited options of cleaning and/or reuse.^40^ Indeed, using HEPA filters on a global scale to combat the transmission of airborne viruses in indoor spaces incur prohibitive environmental and financial costs.

In this work and using a facile fabrication process, we prepared a granular filtration material composed of amyloid nanofibrils (AF) and iron (Fe) oxyhydroxides nanoparticles, i.e., AF-Fe (**Fig. 1a**). The AF were prepared from whey protein extract, a by-product of the dairy industry by lowering the pH to 2 and cooking at 90 °C for ≈5 h. Afterwards, Fe nanoparticles are precipitated on the fibrils by adding FeCl^3^·6H^2^0 and raising the pH to 7. The material is then converted to a granular form by decanting the water from the material and baking it at 80-90°C. The material is then passed through a 12 US mesh sieve (pore size ≈1.7 mm) to remove larger pieces. However, larger aggregates of AF-Fe might form during packaging and storage (**Fig. 1b)**. The material, thus, has a broad size distribution with 50% of its mass smaller than 3 mm, as determined using sieve analysis (**Fig. 1c). Fig. 1d** shows scanning electron microscopy (SEM) images of the material at various magnification scales where the iron oxyhydroxides nanoparticles can be visualized at the highest magnifications. The chemical composition of the material was further verified using Fourier transform infrared spectroscopy (FTIR; **Fig. 1e**), which shows the three peaks for amide groups, representative of the amyloid fibrils, and one of the Fe-O-H group, representative of the iron oxyhydroxides nanoparticles. The material has a high specific surface area, 44.1 m^2^/g (**Fig. 1f**), i.e., the 50% surface coverage capacity of 1 g of the material is ≈ 7 * 10^14^ and ≈ 3 * 10^13^ for 30 or 150 nm virus particles, respectively. Its specific density, ρ^s^, is 2.1 g/cm3 and bulk densities are 1.7 and 1.4 g/cm3 of air-equilibrated and oven-dried samples, respectively, showing a relatively high intra-particle porosity of 36% with 30% volumetric water content. The intra-particle pore-size distribution was further investigated using mercury intrusion porosimetry, revealing pores in size ranges of tens and thousands of nanometers (**Fig 1g, S1**). These pores are in the size range which could serve as trapping cavities for viruses, preventing their release once they are attached to the surface of the AF-Fe.^41^ Collectively, the properties of the granular AF-Fe make it a highly promising candidate to filter virus-loaded aerosols.

**Figure 1.**
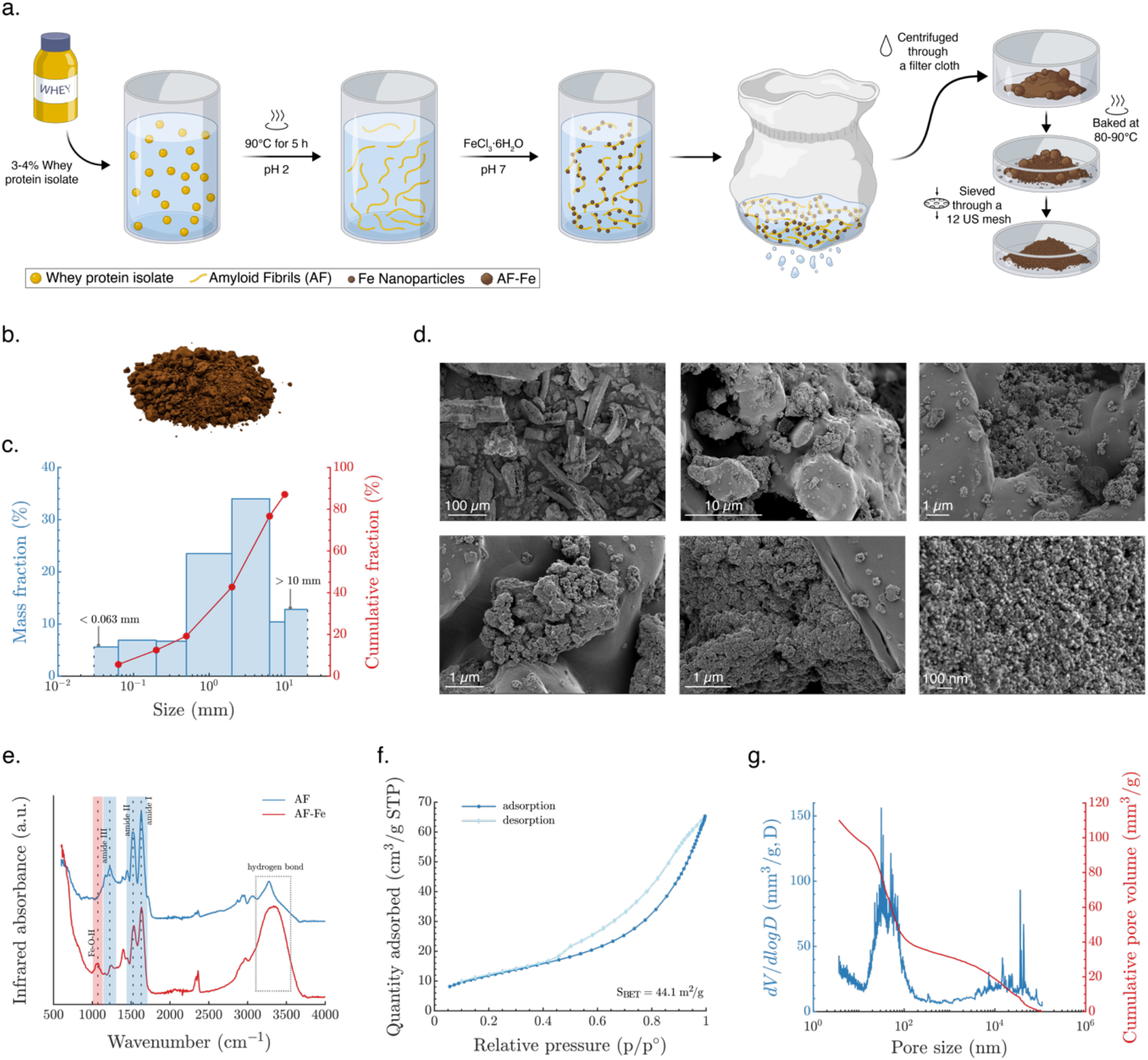
Fabrication and characterization of the granular AF-Fe. (a) A schematic showing the fabrication process of the AF-Fe. (b) A vectorized photo of the material after removing the background. (c) Size distribution of the AF-Fe determined using sieve analysis. (d) SEM images of the material at varying magnifications. (e) FTIR analysis of both AF and AF-Fe. (f) N_2_ adsorption analysis of the AF-Fe to determine its surface area. (g) Intra-particle pore size distribution of the AF-Fe determined using mercury intrusion porosimetry.

To test the filtration efficiency of the material, we designed and built a compact experimental setup (**Fig. 2, S2**) in which virus-loaded aerosols are generated, passed through the AF-Fe at a flowrate of 7.5 l/min, and then collected on a gelatine membrane that traps ≥99% of viruses passing through while maintaining their infectivity (**Fig. S3**). These gelatine membranes are water-soluble, facilitating virus extraction and subsequent infectivity and genome count assessment. The filtration efficiency of AF-Fe was determined by comparing the infectious viruses trapped on the gelatine membranes in the presence of AF-Fe versus in its absence. The setup is compact, allowing for complete operation inside of laminar flow hoods (**Fig. S2**). Additionally, all of its connections are tightly sealed, and its outlet tube is supplemented with an impinger and a HEPA filter to ensure safe operation. A detailed account of the setup and the experimental procedure is presented in the Materials and Methods section.

**Figure 2.**
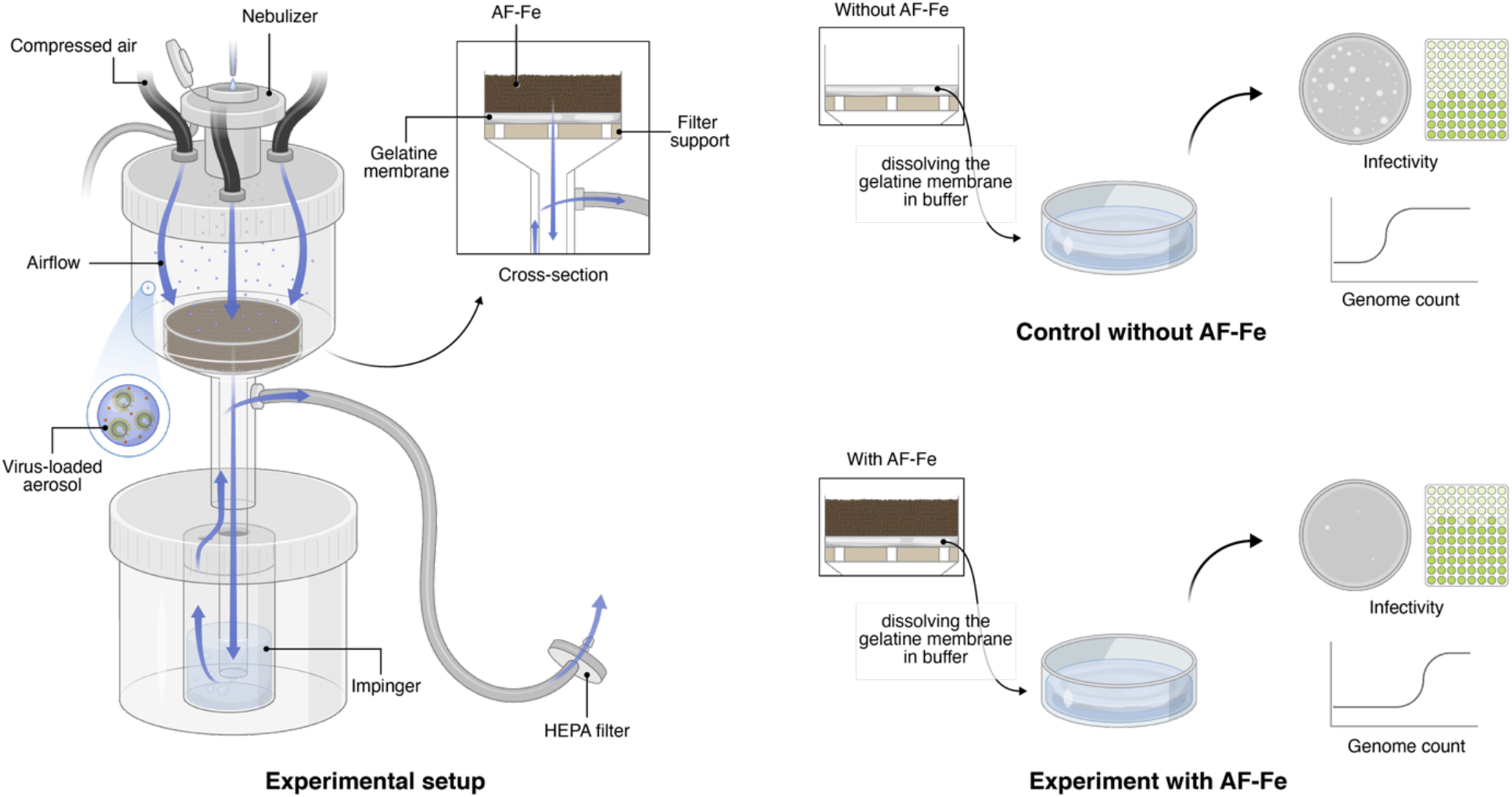
Schematic of the experimental setup and procedure. The experimental setup, left, is composed of: a filter holder with a nebulizer fitted on its top and three hoses connected to three compressed air tanks; the granular AF-Fe placed on a gelatine membrane, which traps the virus-loaded aerosols that pass through the AF-Fe media; an impinger serving as an additional trapping mechanism for any aerosols that go through the gelatine membrane; a HEPA filter as a third safety measure to avoid releasing any virus-loaded aerosols into the surrounding air. For both experiments with (bottom right) and without AF-Fe (top right), the gelatine membrane was dissolved in a buffer to determine the concentration of infectious viruses and the genome count.

The filtration efficiencies of the AF-Fe against H1N1 (the influenza virus strain responsible for the flu pandemic in 2009) and SARS-CoV-2 were, in average, 99.87%and 95.91%, respectively (**Fig. 3a**); both viruses are enveloped and known to be airborne. The filtration efficiency against the enterovirus EV71 was equal to 99.0% (**Fig. 3b**); EV71 is non-enveloped, and is known for its stability and resistance to harsh chemical conditions. Due to safety concerns, most aerosol studies are conducted using bacteriophages.^42^ To situate our results in the context of the existing literature, we tested the efficiency of the AF-Fe against two of the most commonly used bacteriophages in aerosol studies: Φ6, an enveloped bacteriophage that infects *Pseudomonas syringae*; and MS2, a non-enveloped virus that infects *Escherichia coli*. The filtration average efficiency of Φ6 was found to be 99.99% and that of MS2 98.29% (**Fig. 3c**). The latter is known to be one of the most stable viruses in the aerosol phase, and was estimated to be seven times more stable than coronaviruses.^42^ Remarkably, this filtration efficiency is accompanied by a very low pressure-drop across the material, < 0.03 bar (**Fig. S4**), indicating that the AF-Fe would require minimal energy for operation. We further tested the effect of reducing the amount of AF-Fe on both the filtration efficiencies and the pressure-drop. Using as low as two-thirds of the material used in the reported experiments had little to no effect on filtration efficiencies (**Fig. S5, S6**), while reducing the pressure drop to ≈ 0.02 bar (**Fig. S4**).

**Figure 3.**
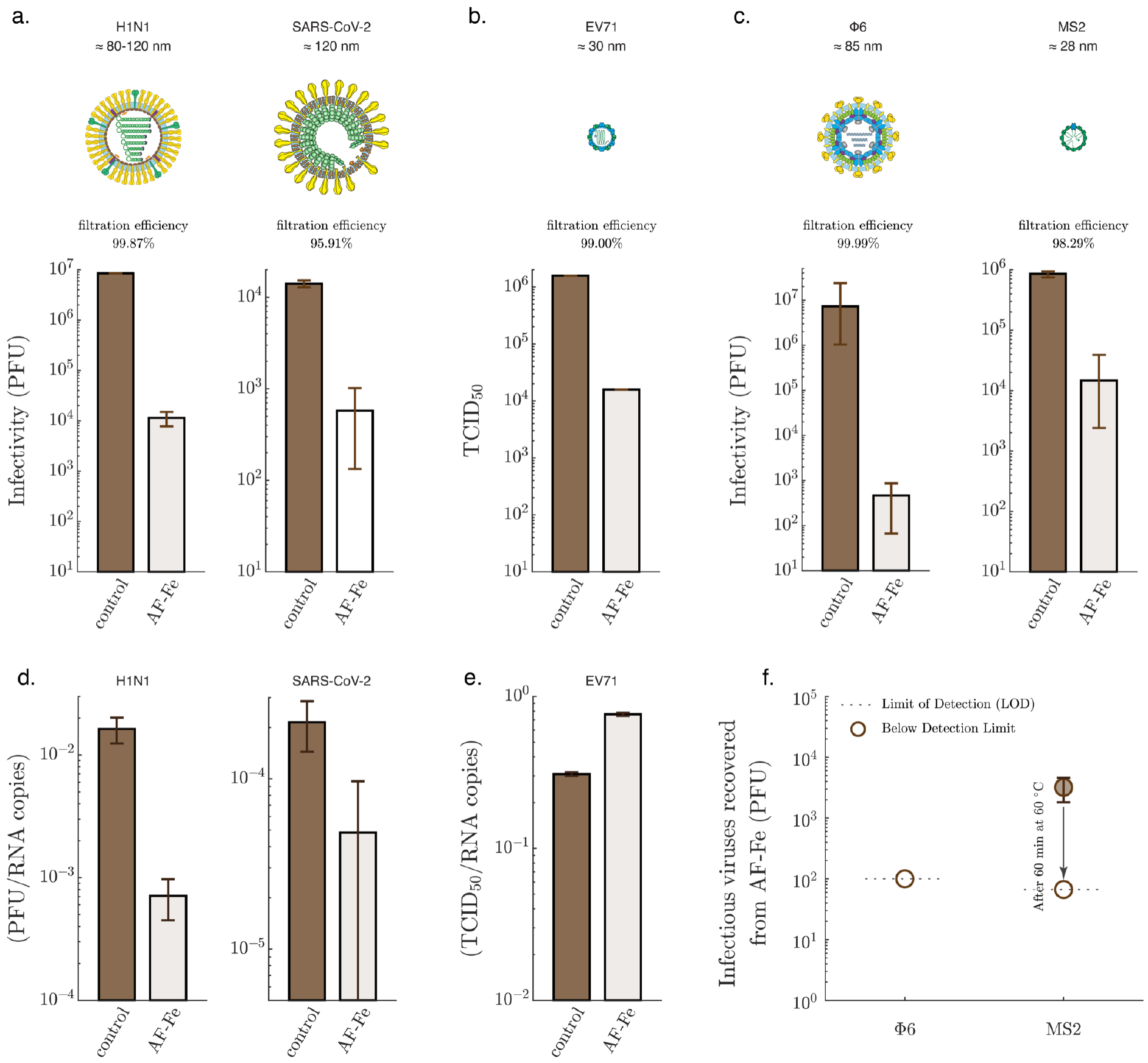
Filtration of virus-loaded aerosols using AF-Fe. Infectious viruses trapped on the gelatine membranes in the absence (control) and the presence of AF-Fe for H1N1 and SARS-CoV-2 (a); EV71 (b); Φ6 and MS2 (c). The plotted values for H1N1 (control and AF-Fe), SARS-CoV-2 (control and AF-Fe), EV71 (control and AF-Fe), and Φ6 (AF-Fe) represent the average from two replicas, with the error bars representing the range. For Φ6 (control) and MS2 (control), the plotted values represent the average from four replicas, with the error bars representing the range. For MS2 (AF-Fe), the plotted value represents the average from three replicas, with the error bar representing the range. For H1N1, SARS-CoV-2, Φ6, and MS2, the infectivity was determined using plaque-forming units (PFU) assays. The infectivity of EV71 was determined using median tissue culture infectious dose (TCID_50_). The ratio of infectious viruses to RNA copies, determined using RT-qPCR, is shown for H1N1 (d), SARS-CoV-2 (d), and EV71 (e). (f) Infectious viruses recovered after incubating the AF-Fe in PBS buffer for ≈1 h for both Φ6 and MS2. Representations of virions were reproduced with permission.^44^

**Fig. 3d** shows that the ratio of infectious H1N1 and SARS-CoV-2 to total genome count decreased after passing the AF-Fe, indicating that the viruses are not just trapped, but are also partially inactivated. A direct interaction between the viruses and AF-Fe would be needed for inactivation to occur. Part of the trapped aerosols is likely to be re-aerosolized again by the airflow shear forces. In this transient period of being attached to the AF-Fe, the viruses are inactivated. However, it also remains possible that the mechanical stress of re-aerosolization contributes to the observed inactivation. Such inactivation, however, was not observed for EV71 (**Fig. 3e**), confirming that non-enveloped viruses are more robust and resistant to mechanical stresses due the interactions with AF-Fe and the re-aerosolization process.

One of the critical challenges presented by commercially-available aerosol filters is handling and recycling contaminated filters. To assess the safe handling of the material, we incubated the AF-Fe for ≈1 h in phosphate-buffered saline (PBS) buffer after filtering aerosols loaded with Φ6 or MS2. No infectious viruses were recovered in the case of Φ6 (**Fig. 3f**), indicating that the AF-Fe completely inactivated the virus and/or trapped it irreversibly. In the case of the non-enveloped MS2, < 0.5% of the infectious viruses were recovered, showing a remarkable capacity for irreversibly trapping the virus. Baking the AF-Fe at 60 °C for 1 h was sufficient to completely inactivate MS2 to below the detection limit (**Fig. 3f**). The inactivating and trapping capacity of AF-Fe is in agreement with previous work, where we show that a membrane composed of AF and iron oxyhydroxides nanoparticles could completely eliminate viruses from bulk water.^31^

To gain deeper insights into the mechanisms of aerosol entrapment by the AF-Fe, we modeled four of the key aerosol entrapment mechanisms (**Fig. 4a**): (i) diffusion, which is driven by Brownian motion of the aerosol droplets; (ii) interception, which occurs when the airflow line comes within one aerosol radius distance from the grains of the filter; (iii) gravitational settling, which is driven by gravitational forces acting on the aerosol particles; and (iv) impaction, which is driven by the inertia of the aerosol particles. All of these processes depend on both the size of the AF-Fe grains, *d*_g_, and the size of the aerosol droplets, *d*_a_. **Fig. 4b** (first four panels) shows that filtration efficiency due to diffusion decreases with increasing *d*_a_; whereas, the efficiencies of interception, gravitational settling, and impaction increase with increasing *d*_a_. The relative contributions of these mechanisms varied considerably with varying the size of grains. While these models were constructed for single-size grain filters, our AF-Fe filter has a very broad size distribution (**Fig. 1c, S9**). To obtain more specific insights into our filter, we calculated an equivalent grain size, *d*_g_ = 0.139mm, satisfying Ergun’s equation (**Eq. S29**),^43^ using the measured pressure drop across the AF-Fe filter, 0.029 bar, as an input parameter (**Fig. 4b;** last panel). Moreover, while the nominal size of aerosol produced by the nebulizer is 4 − 6 μm, we observed that many aerosols were between 0.1 − 1.0 μm in diameter at relative humidity, RH, between 80% – 85% (**Fig. S7**). We, therefore, consider the whole aerosol size range between 0.1 − 10 μm in our discussion. **Fig. 4b and S8** suggest that diffusion and interception are the two key mechanisms for trapping aerosols ≲ 2 μm. For aerosol ≳ 2 μm, all three mechanisms, i.e., interception, gravitational settling, and impaction, exhibit very high efficiencies. The filtration efficiency curve presented in **Fig. 4b** (last panel) resembles that of fiber-based filters, with diffusion, interception, and impaction being considered the key filtration mechanisms.^40^ Our results indicate the AF-Fe can have similar aerosol entrapment mechanisms to fiber-based filters. It is worth mentioning, however, that our model neither considers the effect of AF-Fe polydispersity nor contributions from other factors, such as the irregular shape, roughness, charge, or hydrophilicity of AF-Fe. Future work on developing intricate models that consider the effect of these parameters would be necessary to obtain a more detailed and quantitative assessment of the mechanisms of aerosol filtration using grain-based filters.

**Figure 4.**
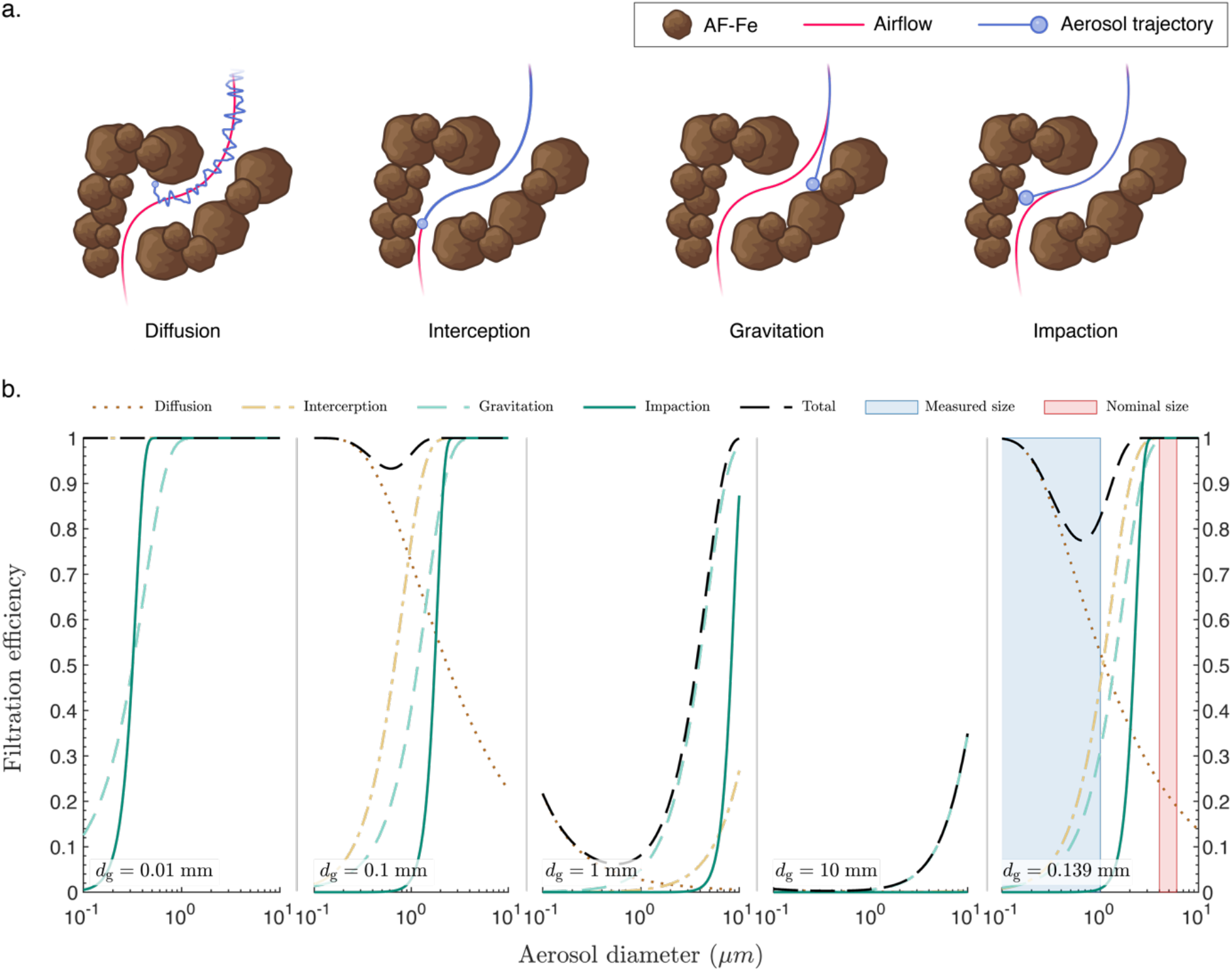
Mechanisms of aerosol entrapment. (a) Schematics depicting different processes involved in aerosol entrapment. (b) Filtration efficiencies due to diffusion, interception, gravitational settling, and impaction through grain bed filters with monodispersed grain size, *d*_g_, which was varied systematically between 0.01 and 10 mm. *d*_g_ = 0.139mm was calculated using Ergun’s equation based on the measured pressure drop across the AF-Fe filter. In (b), the nominal size indicates the diameter range of freshly nebulized droplets according to the manufacturer, while the measured size indicates the range of aerosol diameters at a relative humidity (RH) of 80% − 85% (**Fig. S7**).

Altogether, our results demonstrate that AF-Fe can filter virus-loaded aerosols at high efficiencies as well as in a sustainable and environmentally-friendly manner. The pressure drop across the material is remarkably low, which indicates low energy and cost of operation. In addition, the contaminated material is safe to handle, with markedly more possibilities for recycling than currently-available commercial filters. We foresee the material being used to combat the spread of airborne viruses on a global scale, while exerting virtually no environmental footprint. The use of granular material for aerosol filtration in this study is also expected to inspire the search for new, local, environmentally-friendly materials that could also be used as the main building block for aerosol filters.

## Acknowledgments

The authors gratefully acknowledge support by the Swiss National Science Foundation project N° 31CA30_196217. Prof. Martin Loessner (ETH Zurich) is deeply acknowledged for allowing access to his laboratory facilities; Daniel Kiechl, Peter Bigler, Carmen Saez Garcia Wanzenried, and Rasha Aziz (ETH Zurich) for providing excellent technical support; Dr. Michael Plötze from the ClayLab (ETH Zurich) for conducting the size distribution, density, and porosity determination experiments; Dr. Terttaliisa Lind (PSI) for valuable discussions; Sonia Monti for the design of scientific illustrations; Dr. Eleonora Simeoni, Vivianne Padrun, and Dr. Anna Maria Novello (EPF Lausanne) for biosafety support with SARS-CoV-2 experiments. The authors also gratefully acknowledge the support of the Scientific Center for Optical and Electron Microscopy (ScopeM) of the ETH Zurich.

## Author contributions

A.A. coordinated the study; designed the experimental setup for aerosol experiments; conducted all the aerosol experiments with Φ6, MS2, H1N1, and EV71; analyzed the data; compiled the figures; and wrote the manuscript. H.W. (under the supervision of F.S.) conducted the aerosol experiments with SARS-CoV-2 and evaluated its infectivity. P.A.A. (under the supervision of M.A.) conducted the aerodynamics modeling and calculations; performed aerosol size distribution measurments. C.M. and A.C.Z. (under the supervision of C.T.) propagated H1N1 and EV71, and evaluated their infectivity and genome count; evaluated the genome count of SARS-CoV-2. M.P. performed the FTIR and N^2^ adsorption experiments and analyzed their data. C.G. propagated Φ6 and MS2, and evaluated their infectivity for all aerosol experiments. S.H. carried out the SEM imaging. S.B. synthesized and provided the AF-Fe. A.A. and R.M. wrote the manuscript. R.M. designed and directed the study, aquired funds, analysed data and wrote the manuscript. All authors edited and approved the final manuscript.

## Materials and Methods

A full detailed version of the materials and methods is provided in the Supplementary Information. A summary of the most relevant sections is presented here.

### Materials

#### AF-Fe

The granular AF-Fe material was provided by BluAct (Switzerland) and prepared as detailed later.

#### Viruses

SARS-CoV-2 virus hCoV-19/Switzerland/un-2012212272/2020 was a generous gift from Prof. Isabella Eckerle (University Hospital in Geneva, Geneva, Switzerland). The virus was replicated twice in Vero-E6 cells prior to the experiments. Human H1N1 virus A/Netherlands/602/2009 was a generous gift from Prof. Mirco Schmolke (Department of Microbiology and Molecular Medicine, University of Geneva, Geneva, Switzerland) and was propagated in embryonated chicken eggs.^45^ Enterovirus 71 (EV71) was isolated from a clinical specimen in the University Hospital of Geneva in RD cells and propagated in Vero cells.^46^ Φ6 (21518 DSMZ) and MS2 (13767 DSMZ) bacteriophages were purchased from DSMZ culture collection (Germany) and propagated in their host bacterial cells.

### Methods

#### Preparation of the AF-Fe granular material

Amyloid fibrils were prepared from whey protein isolate (BiPro, Agropur, U.S.A.) by lowering the pH to 2.0 and heating at 90 °C for 5 h.^47^ Then, the amyloid fibrils were coated with iron nanoparticles by mixing FeCl^3^·6H^2^0 and adjusting the pH to 7.0 using NaOH.31 The solution was then centrifuged through a filter cloth to decant the water. Afterward, the retentate was dried at 80-90 °C and sieved through a 12 US mesh (pore size of ≈1700 μm) to remove large particles. It is important to note that some of the material formed larger aggregates during storage and packaging. The material is patented and produced by BluAct (Switzerland) and used as received without further treatment.

#### Assessment of aerosol filtration

The filtration efficiency of AF-Fe against virus-loaded aerosols was assessed using a compact experimental setup composed mainly of a polycarbonate filtration holder (Sartorius, Germany) connected to three 1.5 l compressed air tanks (PanGas, Switzerland) and an Aeroneb® Lab nebulizer unit (Kent Scientific, U.S.A.). Virus-containing solutions were prepared and added to the nebulizer: MS2 and Φ6 were prepared in artificial saliva/mucin solution, H1N1 in allantoid fluid, SARS-CoV-2 in high glucose Dulbecco’s Modified Eagle Medium (DMEM, GlutaMAX™) supplemented with 2.5% fetal bovine serum (FBS), and EV71 in 2.5% serum DMEM. For each virus, 100 μl of the virus-containing solution was nebulized, generating aerosols with an average diameter of 4 to 6 μm, as indicated by the manufacturer, in the upper compartment of the filter holder. The generated aerosols were carried through the filtration media with the airflow from the compressed air tanks at 7.5 l/min (3 × 2.5 l/min). The aerosols passing through the AF-Fe media were then trapped using gelatine membranes (Sartorius, Germany). The membranes are water-soluble and designed to trap virus-loaded aerosols while retaining their infectivity. After disassembling the setup, the membranes were dissolved in 10 ml of PBS buffer for downstream analysis of infectivity and genome count of viruses. The efficiency of the AF-Fe media for the filtration of each virus was determined by comparing the infective viruses in the gelatine membrane in the presence versus the absence of the AF-Fe media. All experiments were conducted in a laminar flow hood.

#### Additional methods

Experimental details on the propagation of Φ6, MS2, H1N1, SARS-CoV-2, and EV71 viruses, infectivity assays, and RT-qPCR assays are given in full in the Supporting Information along with details on the materials and solutions used, experimental setup, and procedure for aerosol filtration, determination of AF-Fe size distribution, FTIR, SEM, N^2^ adsorption, mercury intrusion porosimetry, water content and density determination, pressure drop measurements, determination of aerosol size distribution, and modeling aerosol entrapment mechanisms.

## References

1. Servellita, V. et al. Predominance of antibody-resistant SARS-CoV-2 variants in vaccine breakthrough cases from the San Francisco Bay Area, California. Nature Microbiology 7, 277–288 (2022).

2. Mannar, D. et al. SARS-CoV-2 Omicron variant: Antibody evasion and cryo-EM structure of spike protein–ACE2 complex. Science 375, 760–764 (2022).

3. Liu, L. et al. Striking antibody evasion manifested by the Omicron variant of SARS-CoV-2. Nature 602, 676–681 (2021).

4. Colson, P. et al. Culture and identification of a “Deltamicron” SARS-CoV-2 in a three cases cluster in southern France. Journal of Medical Virology (2022) doi:10.1002/jmv.27789.

5. Pulliam, J. R. C. et al. Increased risk of SARS-CoV-2 reinfection associated with emergence of Omicron in South Africa. Science 4947, (2022).

6. Evans, J. P. et al. Neutralizing antibody responses elicited by SARS-CoV-2 mRNA vaccination wane over time and are boosted by breakthrough infection. Science Translational Medicine (2022) doi:10.1126/scitranslmed.abn8057.

7. Thompson, M. G., Natarajan, K., Irving, S. A. & Rowley, E. A. Effectiveness of a Third Dose of mRNA Vaccines Against COVID-19 – Associated Emergency Department and Urgent Care Encounters and Hospitalizations Among Adults During Periods of Delta and Omicron Variant Predominance — VISION Network, 10 States, August 202. Morbidity and Mortality Weekly Report 71, 255–263 (2022).

8. Gupta, R. K. & Topol, E. J. COVID-19 vaccine breakthrough infections. Science 374, 1561–1562 (2021).

9. Levine-tiefenbrun, M. et al. Waning of SARS-CoV-2 booster viral-load reduction effectiveness. Nature Communications 13, 1–4 (2022).

10. Solís Arce, J. S. et al. COVID-19 vaccine acceptance and hesitancy in low- and middle-income countries. Nature Medicine 27, 1385–1394 (2021).

11. Kerr, J. R. et al. Correlates of intended COVID-19 vaccine acceptance across time and countries: Results from a series of cross-sectional surveys. BMJ Open 11, 1–11 (2021).

12. de Figueiredo, A. & Larson, H. J. Exploratory study of the global intent to accept COVID-19 vaccinations. Communications Medicine 1, 1–10 (2021).

13. Khamsi, R. Can the World Make Enough Coronavirus Vaccine? Nature 580, 578–580 (2020).

14. Wouters, O. J. et al. Challenges in ensuring global access to COVID-19 vaccines: production, affordability, allocation, and deployment. The Lancet 397, 1023–1034 (2021).

15. Liu, Y., Salwi, S. & Drolet, B. C. Multivalue ethical framework for fair global allocation of a COVID-19 vaccine. Journal of Medical Ethics 46, 499–501 (2020).

16. Forni, G. et al. COVID-19 vaccines: where we stand and challenges ahead. Cell Death and Differentiation 28, 626–639 (2021).

17. Krammer, F. SARS-CoV-2 vaccines in development. Nature 586, 516–527 (2020).

18. Sparrow, E. et al. Global production capacity of seasonal and pandemic influenza vaccines in 2019. Vaccine 39, 512–520 (2021).

19. Morris, D. H., Rossine, F. W., Plotkin, J. B. & Levin, S. A. Optimal, near-optimal, and robust epidemic control. Communications Physics 4, 1–8 (2021).

20. Perkins, T. A. & España, G. Optimal Control of the COVID-19 Pandemic with Non-pharmaceutical Interventions. Bulletin of Mathematical Biology 82, 1–24 (2020).

21. Li, Y. et al. The temporal association of introducing and lifting non-pharmaceutical interventions with the time-varying reproduction number (R) of SARS-CoV-2: a modelling study across 131 countries. The Lancet Infectious Diseases 21, 193–202 (2021).

22. Flaxman, S. et al. Estimating the effects of non-pharmaceutical interventions on COVID-19 in Europe. Nature 584, 257–261 (2020).

23. Hatchett, R. J., Mecher, C. E. & Lipsitch, M. Public health interventions and epidemic intensity during the 1918 influenza pandemic. Proceedings of the National Academy of Sciences 104, 7582–7587 (2007).

24. Bagheri, G., Thiede, B., Hejazi, B., Schlenczek, O. & Bodenschatz, E. An upper bound on one-to-one exposure to infectious human respiratory particles. Proceedings of the National Academy of Sciences 118, e2110117118 (2021).

25. Samet, J. M. et al. SARS-CoV-2 indoor air transmission is a threat that can be addressed with science. Proceedings of the National Academy of Sciences 118, 1–5 (2021).

26. Pei, G., Taylor, M. & Rim, D. Human exposure to respiratory aerosols in a ventilated room: Effects of ventilation condition, emission mode, and social distancing. Sustainable Cities and Society 73, 103090 (2021).

27. Bazant, M. Z. & Bush, J. W. M. A guideline to limit indoor airborne transmission of COVID-19. Proceedings of the National Academy of Sciences 118, 1–12 (2021).

28. Koelle, K., Martin, M. A., Antia, R., Lopman, B. & Dean, N. E. The changing epidemiology of SARS-CoV-2. Science 375, 1116–1121 (2022).

29. Karim, N. et al. Sustainable personal protective clothing for healthcare applications: A review. ACS Nano 14, 12313–12340 (2020).

30. Dewey, H. M., Jones, J. M., Keating, M. R. & Budhathoki-Uprety, J. Increased Use of Disinfectants During the COVID-19 Pandemic and Its Potential Impacts on Health and Safety. ACS Chemical Health & Safety 29, 27–38 (2022).

31. Palika, A. et al. An antiviral trap made of protein nanofibrils and iron oxyhydroxide nanoparticles. Nature Nanotechnology 16, 918–925 (2021).

32. Collaborators, C.-19 E. M. Estimating excess mortality due to the COVID-19 pandemic: a systematic analysis of COVID-19-related mortality, 2020 – 21. The Lancet 6736, 1–24 (2022).

33. Adam, D. True covid death toll could be more than double official count. Nature 605, 206 (2022).

34. Josephson, A., Kilic, T. & Michler, J. D. Socioeconomic impacts of COVID-19 in low-income countries. Nature Human Behaviour 5, 557–565 (2021).

35. Wang, C. C. et al. Airborne transmission of respiratory viruses. Science 373, eabd9149 (2021).

36. Port, J. R. et al. Increased small particle aerosol transmission of B.1.1.7 compared with SARS-CoV-2 lineage A in vivo. Nature Microbiology 7, 213–223 (2022).

37. Stadnytskyi, V., Bax, C. E., Bax, A. & Anfinrud, P. The airborne lifetime of small speech droplets and their potential importance in SARS-CoV-2 transmission. Proceedings of the National Academy of Sciences 117, 11875–11877 (2020).

38. Lewis, D. Why the WHO took two years to say COVID is airborne. Nature 604, 26–31 (2022).

39. Henning, L. M. et al. Review on Polymeric, Inorganic, and Composite Materials for Air Filters: From Processing to Properties. Advanced Energy and Sustainability Research 2, 2100005 (2021).

40. First, M. W. HEPA FILTERS. Journal of the American BIological Safety Association 3, 33–42 (1998).

41. Canh, V. D. et al. Evaluation of porous carbon adsorbents made from rice husks for virus removal in water. Water (Basel) 13, 1280 (2021).

42. Turgeon, N., Toulouse, M. J., Martel, B., Moineau, S. & Duchaine, C. Comparison of five bacteriophages as models for viral aerosol studies. Applied and Environmental Microbiology 80, 4242–4250 (2014).

43. Ergun, S. Fluid flow through packed columns. Chem Eng Prog 48, 89–94 (1952).

44. Philippe Le Mercier. ViralZone. SIB Swiss Institute of Bioinformatics.

45. Riegger, D. et al. The Nucleoprotein of Newly Emerged H7N9 Influenza A Virus Harbors a Unique Motif Conferring Resistance to Antiviral Human MxA. Journal of Virology 89, 2241–2252 (2015).

46. Tseligka, E. D. et al. A VP1 mutation acquired during an enterovirus 71 disseminated infection confers heparan sulfate binding ability and modulates ex vivo tropism. PLoS Pathogens 14, 1–25 (2018).

47. Jung, J. M., Savin, G., Pouzot, M., Schmitt, C. & Mezzenga, R. Structure of heat-induced β-lactoglobulin aggregates and their complexes with sodium-dodecyl sulfate. Biomacromolecules 9, 2477–2486 (2008).

