## Supplementary information for "Trapping virus-loaded aerosols using granular protein nanofibrils and iron oxyhydroxides nanoparticles"

Table of Contents


| | |
|---|---|
| Supporting Materials and Methods | Page 3-10 |
| Supplementary Figure S1 | Page 14 |
| Supplementary Figure S2 | Page 15 |
| Supplementary Figure S3 | Page 16 |
| Supplementary Figure S4 | Page 17 |
| Supplementary Figure S5 | Page 18 |
| Supplementary Figure S6 | Page 19 |
| Supplementary Figure S7 | Page 20 |
| Supplementary Figure S8 | Page 21 |
| Supplementary Figure S9 | Page 22 |

### S1. Materials and Methods

#### *S1.1 Materials*

Tryptone, agar, 0.2 µm syringe filters, polyethylene glycol with average molecular weight of 5000 to 7000 (PEG 6000), sodium hydroxide (NaOH), sodium chloride (NaCl), sodium bicarbonate ($NaHCO_3$), sodium phosphate monobasic dihydrate ($NaH_2PO_4 \cdot 2H_2O$), monobasic potassium phosphate ($KH_2PO_4$), magnesium sulfate ($MgSO_4$), calcium chloride ($CaCl_2$), phosphate-buffered saline tablets (PBS), paraformaldehyde (PFA) 4% solution in PBS, bovine serum albumin (BSA), HEPES, tosylsulfonyl phenylalanyl chloromethyl ketone (TPCK)-treated trypsin, crystal violet, artificial saliva for pharmaceutical research, and mucin from porcine stomach were purchased from Merck (Darmstadt, Germany); Serum-free Dulbecco's Modified Eagle's Medium (DMEM), DMEM GlutaMAX$^{TM}$, and Fetal bovine serum (FBS) from Life Technologies (Carlsbad, U.S.A.); 1% penicillin/streptavidin from ThermoFisher (Waltham, MA U.S.A.), Avicel GP3515 from DuPont (Thailand); EZNA viral extraction kit from Omega Biotek (Norcross, U.S.A.); QuantiTect kit (#204443) from Qiagen (Hilden, Germany); TPCK Treated from Worthington (Lakewood, U.S.A.); yeast extract from LLG GmbH (Meckenheim, Germany). Polycarbonate filtration holders and gelatine membranes for environmental air monitoring from Sartorius (Germany). Cytiva Whatman™ HEPA-vent filter from Whatman (U.S.A.). SintoGard from Wetrok (Switzerland). Javel 13-14% from Reactolab (Switzerland). Virkon S from Lanxess (Germany). Compressed air tanks, compressed air tubing, and worm screw clamp from PanGas (Switzerland). Φ6 bacteriophage (DSM no. 21518), *Pseudomonas sp.* (DSM no. 21482), MS2 bacteriophage (DSM no. 13767), and *Escherichia coli* (*E. coli*; DSM no. 5695) were purchased from DSMZ-German Collection of Microorganisms and Cell Cultures GmbH (Braunschweig, Germany). MDCK (ATCC CCL-34) cells and Vero 76 (ATCC CRL-1587) were purchased from the American Type Culture Collection (ATCC, Manassas, USA). ). Vero E6 cells (Vero C1008, ATCC CRL-1586) were kindly provided by Prof. Gary Kobinger. H1N1 virus A/Netherlands/602/2009 was a generous gift from Prof. Mirco Schmolke (Department of Microbiology and Molecular medicine, University of Geneva, Geneva, Switzerland). SARS-CoV-2 virus hCoV-19/Switzerland/un-2012212272/2020 (EPI_ISL_2131446) was a generous gift from Prof. Isabella Eckerle (University Hospital of Geneva, Geneva, Switzerland). Enterovirus 71 (EV71) was isolated from a clinical specimen in the University Hospital of Geneva (Geneva, Switzerland).

#### *S1.2 Solutions*

***Artificial Saliva/Mucin solution.*** A solution composed of artificial saliva and 0.3% mucin from porcine stomach was used to mimic respiratory droplets.[1] Approximately 30 mg of mucin was dissolved in 10 ml of artificial saliva by vigorous mixing and sonication.

***Phosphate-buffered Saline (PBS buffer).*** PBS buffer was prepared by dissolving one PBS tablet per 200 ml of Milli-Q water. The buffer was then autoclaved and stored at room temperature until use.

***Luria Bertani (LB) solution.*** LB solution was prepared by dissolving 5 g/L of NaCl, 10 g/L of tryptone and 5 g/L of yeast extract in 1 L of MilliQ-water followed by autoclaving.

***LB Agar.*** LB agar was prepared by adding 15 g/L of agar to Luria Bertani (LB) solution. After autoclaving, the still warm solution was used to prepare LB agar petri dishes.

***LC Soft Agar.*** LC soft agar was prepared by dissolving 7.5 g/L of NaCl, 10 g/L of tryptone, 5 g/L of yeast extract, 10 g/L of glucose, and 4 g/L of agar to 1 L of MilliQ-water. After autoclaving, 2 mM $MgSO_4$ and 10 mM $CaCl_2$ were added from 0.22 µM filter sterile solutions, followed by aliquoting into test tubes. LC soft agar was used to determine the plaque-forming units (PFU) for both Φ6 and MS2 using the soft agar overlay method.[2]

***Virus Dilution Buffer (VDB).*** VDB contained 0.78 g/l $NaH_2PO_4{\cdot}2H_2O$ and 0.58 g/l NaCl and had a pH of 7.4, adjusted using 1M NaOH. VDB was used for MS2 stocks.

***Buffer A.*** Buffer A was prepared by adding 10 mmol/l $KH_2PO_4$ and 1 mmol/l $MgSO_4$ to Milli-Q water and adjusting the final pH to 7.5 using NaOH. Buffer A was used for MS2 stocks.

#### *S1.3 Cells and viruses*

**Cell lines.** MDCK (ATCC CCL-34) cells, Vero 76 (ATCC CRL-1587) and Vero E6 (ATCC CRL-1586) were cultured in DMEM supplemented with GlutaMAX$^{TM}$, Sodium Pyruvate, Phenol Red, 10% fetal bovine serum (FBS), and 1% Penicillin-Streptomycin and grown at 37˚C in an atmosphere of 5% $CO_2$.

**H1N1.** H1N1 was propagated in embryonated chicken eggs according to Brauer and Chen (2015).[5] Briefly: Serum pathogen-free fertilized chicken eggs were incubated at 37 °C and 55-60% humidity for 11 days. The virus inoculation was then carried out by injecting 100 FPU stock into the allantoic cavity using a needle. After 2 days of incubation at 37 °C, the eggs were chilled for at least 24 h at 4 °C. The eggshell above the air sac and the chorioallantoic membrane was then opened, and the allantoic fluid containing the virus was harvested. The fluid was cleared from debris by centrifugation, aliquoted, and transferred to -80 °C for long-term storage. Viral titers, assessed by plaque assay in MDCK cells, were ≈ $10^9$ PFU/ml.

**SARS-CoV-2.** hCoV-19/Switzerland/un-2012212272/2020 was used for the filtration experiments. The virus was propagated in Vero E6 cells; the supernatant was collected 3 days post-infection, clarified, aliquoted, and frozen at −80°C, and subsequently titrated by plaque assay in Vero-E6 cells (titer ≈ $3 \times 10^6$ PFU/ml).

**Enterovirus 71 (EV71).** EV71 was amplified in Vero cells. For viral stock production, the cells were infected in DMEM + 2.5% FBS for 1 hour at 37˚C. The inoculum was then removed, and a fresh medium was added. The infectious supernatant was collected 72 h post-infection, aliquoted, and frozen at -80˚C before titration. The virus was titrated using median tissue culture infectious dose ($TCID_{50}$).

**Φ6.** Φ6 (DSM no. 21518) was propagated and purified according to Block et al. (2014)[3] and Palika et al. (2021)[4]. Briefly: Φ6 was harvested from soft agar plate lysate of inoculated host bacteria, *Pseudomonas sp.* (DSM no. 21482), after incubation at room temperature overnight. Cell debris and agar were removed by centrifugation. Φ6 was then salted out in 0.5 M NaCl and collected on a PEG pellet by centrifugation. After resuspending in Buffer A the PEG was removed by centrifugation. The phage was then purified in three ultracentrifugation steps: one in buffer and two in sucrose gradients. The purified phage had a final concentration of ≈ $10^{12}$ PFU/ml.

**MS2.** MS2 (DSM no. 13767) was harvested from soft agar plate lysate of inoculated host bacteria, *E. coli* (DSM np. 5695), after incubation at 37˚C overnight. Cell debris and agar were removed by filtering through 0.2 µm syringe filters followed by centrifugation. The phage stock was stored in VDB buffer and had a final concentration of ≈ $10^8$ PFU/ml.

#### *S1.5 Infectivity assays*

**H1N1.** The infectivity of H1N1 was determined using a plaque assay. Confluent cultures of MDCK cells in 6 multiwell plates were incubated at 37˚C for 1 h with 10-fold serial dilutions of the virus, prepared in serum-free DMEM containing 1% penicillin-streptomycin. Upon inoculum removal, the cells were washed and overlaid with MEM containing 0.3% BSA, 0.9% Bacto agar, and 1 µg/mL TPCK-treated trypsin. After 48 h of incubation at 37˚C, the cells were fixed with 4% formaldehyde solution and then stained with 0.1% crystal violet. The number of PFUs per dilution was determined using a fine-scale magnifying comparator and a white light table.

**EV71.** The infectivity of EV71 was determined by $TCID_{50}$. Confluent cultures of Vero cells in 96 multiwell plates were incubated at 37˚C for several days with 10-fold serial dilutions of the virus prepared in DMEM containing 2% FBS and 1% penicillin-streptomycin. Each sample was analyzed in duplicate. The cytopathic effect was checked every day, and the final analysis was performed when there was no more variation compared to the previous day. The infectious virus concentrations were thus calculated according to Spearman[6] and Kärber[7] as described by Hierholzer and Killington (1996)[8]. The final virus concentration was expressed as $TCID_{50}$/ml.

**SARS-CoV-2.** The infectivity of SARS-CoV-2 was determined using a plaque assay. Confluent cultures of Vero-E6 cells in 24-well plates were incubated at 37 ˚C for 1 h, with 5-fold serial dilutions of the virus prepared in DMEM GlutaMAX™ supplemented with 2.5% FBS and 1% penicillin-streptomycin. Then, the viral inoculum was removed, and cells were overlaid with DMEM containing 2.5% FBS, 1% penicillin-streptomycin and 0.4% Avicel GP3515. After 48 h of incubation at 37 ˚C, the cells were fixed with 4% PFA and stained with 0.1% crystal violet. The number of PFUs were counted and final virus concentration was expressed as PFU/mL. Each samples wad analyzed in triplicates (for the 1:1 and 1:5 dilutions, 5 replicas were performed).

**Φ6.** The infectivity of Φ6 was determined using the soft agar overlay method using 10-fold serial dilutions. Each dilution was assessed using three technical replicas. For each plate, 10 µL Φ6-containing solution was mixed with 200 µL of log-phase growing host bacteria, *Pseudomonas sp.* (DSM no. 21482), in 5 ml of LC soft agar solution. The mixture was then spread on a LB agar plate and incubated overnight at room temperature. The infectious virus concentrations were thus expressed as PFU/ml.

**MS2.** The infectivity of MS2 was determined by the soft agar overlay method using 10-fold serial dilutions. Each dilution was assessed using three technical replicas. For each plate, 100 µL MS2-containing solution was mixed with 100 µL of log-phase growing host bacteria, *E. coli* (DSM np. 5695), in 5 ml of LC soft agar solution. The mixture was then spread on a LB agar plate and incubated overnight at 37 ˚C. The infectious virus concentrations were thus expressed as PFU/ml.

#### *S1.6 Genome count using reverse transcriptase real-time polymerase chain reaction (RT-qPCR)*

**H1N1, SARS-CoV-2, and EV71.** EZNA viral extraction kit (Omega Biotek) was used to extract the viral genome and quantified using RT-qPCR with the QuantiTect kit (#204443; Qiagen, Hilden, Germany) in a StepOne ABI Thermocycler. Using the slope-intercept form, $C_t$ values were converted into RNA concentration. The $R^2$ for all experiments ranged between 0.96-0.99. Statistical analysis was done with Prism software (Prism 8, GraphPad). Primers used for each virus are listed in **Table S1**. Four ten-fold dilution series of *in vitro* transcripts of the influenza

A/California/7/2009(H1N1) M gene, SARS-CoV-2 E gene, and EV71 VP1 gene were used as reference standards.

**Table S1.** Primers used for the RT-qPCR analysis of H1N1, SARS-CoV-2, and EV71.

| Primer | Sequence (5'-3') | Target |
|---|---|---|
| H1N1-F | GACCRATCCTGTCACCTCTGAC | M gene |
| H1N1-R | AGGGCATTYTGGACAAAKCGTCTA | |
| H1N1-probe | TGCAGTCCTCGCTCACTG GGCACG | |
| EV71-F | GCCAGATTCCAGAGGGTCTCTCGCATGGC | VP1 gene |
| EV71-R | GCCATGCGAGAGACCCTCTGGAATCTGGC | |
| EV71-probe | GCGGAACCGACTACTTTGGG | |
| SARS-CoV-2-F | ACA-GGT-ACG-TTA-ATA-GTT-AAT-AGC-GT | E gene |
| SARS-CoV-2-R | ATA-TTG-CAG-CAG-TAC-GCA-CAC-A | |
| SARS-CoV-2-probe | ACA-CTA-GCC-ATC-CTT-ACT-GCG-CTT-CG | |

#### *S1.7 Experimental setup assembly and operation*

**Setup assembly.** The filtration setup and the nebulizer unit were sterilized before each experiment either by thorough treatment with SintoGard (for experiments with bacteriophages) or autoclaving (for experiments with human viruses). The walls of the upper chamber and the nebulizer head were then treated with an anti-fogging coat (Zeiss, Germany) to minimize loss of aerosol by condensation; applying the anti-fogging coat was not done for experiments conducted in biosafety level 3 laboratories. The impinger was filled with PBS for MS2 and Φ6 experiments, Javel 13-14% (Reactolab, Switzerland) for H1N1 experiments, water aspirated at the end of the experiments with Virkon S (Lanxess, Germany) for SARS-CoV-2 experiments, and SintoGard (Wetrok, Switzerland) for EV71 experiments. The setup was then assembled bottom-up. The gelatine membrane was added on the top of the filter support and fixed in place by screwing the upper chamber tight while making sure not to crack the membrane. The desired amount of AF-Fe is then added to the top of the gelatine membrane. The upper cap of the setup was screwed tightly and connected to the compressed air tanks and the nebulizer head. For experiments with human viruses, the outlet of the setup was complemented with a Cytiva HEPA-Vent Filter (Whatman, U.S.A.), and all the connections were additionally sealed using parafilm. Experiments with H1N1, EV71, Φ6, and MS2 were conducted in biosafety level 2 laboratories; experiments with SARS-CoV-2 were conducted in biosafety level 3 laboratories.

**Assessing the efficiency of gelatine membranes.** According to the manufacturer, the gelatin membranes traps viruses in the aerosol phase with an efficiency of 99.94%. This efficiency might change based on the aerosol size and the viruses. The efficiency of the gelatine membranes was tested by comparing the infectious viruses trapped in the membranes to that in the PBS buffer of the impinger for both Φ6 and MS2.

**Assessing the infectivity of viruses retained on the AF-Fe.** Infectious viruses trapped on the AF-Fe media were assessed by incubating the material for ≈1h in PBS buffer and running infectivity assays on the supernatant. This was only possible for bacteriophages due to the cytotoxic effects of the AF-Fe on the host cells of the human viruses.

#### *S1.8 AF-Fe characterization*

**Size distributions**. Size distribution of the AF-Fe was determined using a combined sieve analysis and laser scattering techniques. Sieve analysis was conducted using a vibratory sieve Shaker AS 200 Control from Retsch (Germany). The analysis was done according to the standard DIN IS0 3310-3 with cut-off sizes 0.063, 0.2, 0.5, 2.0, 6.3, and 10.0 mm. The sieving continued for 3 min with 1 min intervals and amplitude of 1 mm. A total of ≈ 110 g of air-dried material was used with a material loss of 1.7 %. The sieves were 200 mm in diameter and 50 mm in height. The fraction that went through the 0.063 mm cut-off has been further characterized after dispersing in water (0.1% Na-hexametaphosphate solution) using LA-950 Laser Scattering Particle Size Distribution Analyzer from Horiba (Japan). Data were analysed using the EasySieve Comfort v2.4 (Retsch) and LA-950 NextGen v8.5 (Horiba) software for the sieve and laser scattering measurement, respectively.

**Scanning electron microscopy (SEM).** Fractions of different sizes of the material were mounted on SEM aluminum pin stubs with double adhesive carbon tape or with conductive carbon glue (Plano GmbH, Germany) for larger pieces. After drying, the samples were sputter-coated with 4 nm of platinum/palladium (CCU-10, Safematic, Switzerland). SE-inlens and Everhart-Thornley (ET) SE-images were recorded at a working distance of around 4-5 mm with a scanning electron microscope (Merlin, Zeiss, DE), operated at an accelerating voltage of 1.5 kV.

**Fourier transform infrared spectroscopy (FTIR).** FTIR was carried out on pure AF and AF-Fe. The samples were scanned at room temperature over a wavelength range of 600-4000 $cm^{-1}$ at a resolution of 2 $cm^{-1}$ using a Varian 640 spectrometer (Agilent Technologies, USA) equipped with a Golden Gate diamond ATR stage.

**$N_2$ adsorption analysis.** The specific surface area and pore volume of AF-Fe were determined by $N_2$ adsorption analysis at -196 °C with TriStar II (Micromeritics, U.S.A.). The visible area was calculated according to the Brunauer-Emmett-Teller (BET) method. The total pore volume was determined based on the amount of adsorbed nitrogen at $p/p_o$ = 0.96. Additionally, the pore size distribution was calculated with Barrett-Joyner-Halenda's (BJH) pore size distribution, assuming slit-shaped pores using Micromeritics 3Flex (Micromeritics, U.S.A.). The samples were degassed under $N_2$ at 150 °C for 24 h before measurements.

**Water Content, density, and porosity.** Water content ($u$) with respect to the dry mass of the material mass was determined by measuring the mass of a sample before and after drying at 105 ˚C overnight until weight constancy. Specific density ($\rho_s$) was determined using a gas displacement system AccuPycTM 1340 (Micromeritics, U.S.A.) of the 105 °C dried fine-grained sample. Bulk density ($\rho$) was determined using an envelope density analyzer GeoPycR 1305 (Micromeritics, U.S.A.) of an air-dried large piece sample (≈1 cm in diameter). Density measurements were conducted in duplicates (each an average of 10 measurements). The specific density, bulk density, and water content were used to determine the volumetric, dry bulk density ($\rho_{\mathrm{dry}}$), water content ($\theta$), and total intra-particle porosity ($n$) according to the following equations.

$$\rho_{\mathrm{dry}} = \frac{\rho}{1+u} \qquad \textbf{Eq. S1}$$

$$\theta = \frac{u}{\rho_{\mathrm{dry}}} * 100 \qquad \textbf{Eq. S2}$$

$$n = 1 - \frac{\rho_{\mathrm{dry}}}{\rho_{\mathrm{s}}} \qquad \text{Eq. S3}$$

**Mercury intrusion porosimetry (MIP).** MIP was used to determine the percentage of open pores and their sizes that are Hg-accessible. MIP was carried out with a combined instrument (Pascal 140 + 440, POROTEC) to measure macro- and mesopores in the range 1.8 - 58000 nm radius. Samples of the fraction 6.3 – 10 mm were used. First, the air-dried specimens were evacuated for 30 min to dry (<0.03 kPa). Then the measurements were conducted by incrementing the pressure up to 400 MPa on a sample immersed in the non-wetting mercury. Hereby, the rate of pressure increase was automatically adjusted in an advanced procedure with lower rates at lower pressure levels and during measured intrusion processes. With increasing pressure, mercury intrudes into progressively smaller voids. The pore volume can be derived from the quantity of intruded mercury. The pore size distribution can be determined according to the Washburn equation (**Eq. S4**), which gives a relationship between pressure and pore size.[9]

$$r = \frac{2\gamma \cos\theta_{\mathrm{Hg}}}{p}, \qquad \text{Eq. S4}$$

where $r$ is the pore radius, $p$ pressure, $\gamma$ surface tension of mercury (0.48 N/m), and $\theta_{\mathrm{Hg}}$ the wetting angle of mercury (140°).

#### *S1.9 Pressure drop measurement*

The pressure drop was measured under an airflow rate of 8 l/min using a digital manometer Leo1 from Keller (Switzerland). The measurements were conducted using the same setup for running the experiments with virus-loaded aerosols after detaching the lower chamber, the impinger, and the outlet tube with the HEPA filter. The manometer was connected to the upper chamber to record the pressure. The pressure in the presence of the gelatine membrane only was recorded (0.9735 ± 0.0005 bar; error reflects the fluctuation over the measurement period) and used as a reference reading. AF-Fe was added in increments of 8 g, and the pressure inside the upper chamber was recorded and subtracted from the pressure in the presence of the gelatine membrane only.

#### *S1.10 Inter-particle porosity*

The inter-particle porosity was estimated based on the volume occupied by the AF-Fe and its bulk density. The volume ($V_T$) per 8 g of the material was estimated using 15 ml graduated centrifuge tubes; the volume ranged between 7 and 9 ml. The porosity (φ) was thus calculated according to the following formula (with $m$ being the mass of AF-Fe, i.e., 8 g).

$$\varphi = \frac{V_T - m/\rho}{V_T} \qquad \text{Eq. S5}$$

This yielded $\varphi = 30 - 40\%$, approximated to 10% steps.

#### *S1.11 Aerosol size distribution*

Aerosol particles size distribution was measured using a scanning mobility particle sizer (SMPS) consisting of a homemade differential mobility analyzer (DMA, 93.5 cm long, 0.937 cm inner diameter and 1.961 outer diameter) and a condensation particle counter (CPC, Model 3775, TSI Inc.). Solutions were nebulized in the upper chamber of the filter setup using the same operating conditions except without any AF-Fe. The SMPS sampled particles from the total airflow at a rate of 0.3 L/min. The relative humidity, RH, in the air was between 80% and 85% measured with an inline humidity probes in the aerosol flow and the DMA sheath flow. The uncertainty in RH was ± 2.5 %. The RH was the result of equilibrating aerosol evaporation into the dry compressed air stream that carried the particles to the SMPS.

#### *S1.12 Modelling aerosol entrapment mechanisms*

Filter efficiency to capture aerosol particles from the air stream was calculated considering four different processes: diffusion, interception, gravitational settling, and impaction according to Otani et al (1989)[10] for a monodisperse grain size distribution. The single grain efficiency due to gravitational settling, $\eta_{\mathrm{G}}$, is determined from

$$\eta_{\mathrm{G}} = \frac{G}{1+G} \qquad \text{Eq. S6}$$

and the gravity parameter, $G$, given by

$$G = \frac{(\rho_{\mathrm{a}} - \rho_{\mathrm{air}}) g C_{\mathrm{c}} {d_a}^2}{18\mu U}, \qquad \text{Eq. S7}$$

where $\rho_{\mathrm{a}} = 1.7$ g/cm$^3$ is the assumed density of aerosol particles, $\rho_{\mathrm{air}} = 1.1 \times 10^{-3}$ g/cm$^3$ is the air density, $g = 9.8$ m/s$^2$ is the acceleration of gravity, $C_{\mathrm{c}}$ is the Cunningham slip correction factor, $d_{\mathrm{a}}$ is the aerosol particle diameter, $\mu = 1.8 \times 10^{-4}$ g/cm$^1$/s$^1$ is the dynamic gas viscosity and $U = 7.2$ cm/s$^1$ is the interstitial air velocity. We calculate $C_{\mathrm{c}}$ for aerosol particles from a previous parameterization[11] using the mean free path of air molecules, $\lambda$, and the Knudsen number, $Kn$, following

$$\lambda = \lambda_0 \left(\frac{P_0}{P}\right)\left(\frac{T}{T_0}\right)\left(\frac{1 + T_{\mathrm{c}}/T_0}{1 + T_{\mathrm{c}}/T}\right), \qquad \text{Eq. S8}$$

where $\lambda_0 = 6.6 \times 10^{-2}$ µm, $T_0 = 293$ K, $T_{\mathrm{c}} = 110$ K, $P_0 = 1013$ mbar, $T$ is the temperature, $P$ is the pressure,

$$Kn = \frac{2\lambda}{d_{\mathrm{a}}}, \qquad \text{Eq. S9}$$

and

$$C_{\mathrm{c}} = 1 + Kn\left(1.142 + 0.558 e^{-0.999/Kn}\right). \qquad \text{Eq. S10}$$

Then, the gravitation filtration efficiency, $E_{\mathrm{G}}$, is determined by

$$E_{\mathrm{G}} = 1 - e^{-\frac{3\eta_{\mathrm{G}}\alpha L}{2(1-\alpha)d_{\mathrm{g}}}}, \qquad \text{Eq. S11}$$

where $\alpha = 0.68$ is the packing density, $L = 2$ cm is the filter thickness, and $d_\mathrm{g}$ is the grain diameter. The single grain efficiency due to diffusion, $\eta_\mathrm{D}$, is a function of the particle diffusion coefficient, $D$, the Reynolds number, $Re$, and the Schmidt number, $Sc$, following

$$D = \frac{k_\mathrm{b} T C_\mathrm{c}}{3\pi\mu d_\mathrm{a}}, \qquad \text{Eq. S12}$$

$$Re = \frac{\rho_\mathrm{air} U d_\mathrm{g}}{\mu}, \qquad \text{Eq. S13}$$

and

$$Sc = \frac{\nu}{D}, \qquad \text{Eq. S14}$$

where $k_\mathrm{b}$ is the Boltzmann constant and $\nu = 1.5 \times 10^{-1}$ $\mathrm{cm^2/s}$ is the kinematic viscosity of air. Then,

$$\eta_\mathrm{D} = a_1 Re^{a_2} Sc^{a_3}, \qquad \text{Eq. S15}$$

where

$$a_1 = 8, a_2 = -2/3 \text{ for } Re < 30$$

$$a_1 = 40, a_2 = -1.15 \text{ for } 30 \le Re < 100 \qquad \text{Eq. S16}$$

$$a_1 = 2.1, a_2 = -1/2 \text{ for } Re \ge 100$$

and

$$a_3 = -\frac{2}{3} + \frac{Re^3}{6(Re^3 + 2 \times 10^5)}. \qquad \text{Eq. S17}$$

Then, the diffusion filtration efficiency, $E_\mathrm{D}$, is determined by

$$E_\mathrm{D} = 1 - e^{-\frac{3\eta_\mathrm{D}\alpha L}{2(1-\alpha)d_\mathrm{g}}}. \qquad \text{Eq. S18}$$

The single grain efficiency due to impaction, $\eta_\mathrm{I}$, is a function of the effective Stokes number, $Stk_\mathrm{eff}$, and the interception parameter, $R_\mathrm{I}$, following

$$Stk_\mathrm{eff} = Stk\left(1 + \frac{1.75 Re(1-\alpha)}{150\alpha}\right) \qquad \text{Eq. S19}$$

where we can write

$$R_\mathrm{I} = \frac{d_\mathrm{a}}{d_\mathrm{g}}, \qquad \text{Eq. S20}$$

and the Stokes number, $Stk$, as

$$Stk = \frac{\rho_\mathrm{p} U C_\mathrm{c} {d_\mathrm{a}}^2}{9 \mu d_\mathrm{g}}. \qquad \text{Eq. S21}$$

Then,

$$\eta_\mathrm{I} = \frac{{Stk_\mathrm{eff}}^3}{1.4 \times 10^{-2} + {Stk_\mathrm{eff}}^3}, \qquad \text{Eq. S22}$$

and the impaction filtration efficiency is

$$E_\mathrm{I} = 1 - e^{-\frac{3\eta_\mathrm{I}\alpha L}{2(1-\alpha)d_\mathrm{g}}}. \qquad \text{Eq. S23}$$

The single grain efficiency due to interception, $\eta_\mathrm{R}$, is a function of $Re$ given by

$$\eta_\mathrm{R} = 16 {R_I}^{a_4}, \qquad \text{Eq. S24}$$

where the exponent is a parameter given by

$$a_4 = 2 - \frac{Re}{(Re^{-1/3} + 1)^3}. \qquad \text{Eq. S25}$$

Then, the interception filtration efficiency is

$$E_\mathrm{R} = 1 - e^{-\frac{3\eta_\mathrm{R}\alpha L}{2(1-\alpha)d_\mathrm{g}}}. \qquad \text{Eq. S26}$$

The total single grain efficiency from all processes, $\eta_\mathrm{tot}$, is

$$\eta_\mathrm{tot} = \eta_\mathrm{G} + \eta_\mathrm{D} + \eta_I + \eta_\mathrm{R}, \qquad \text{Eq. S27}$$

and the total filtration efficiency, $E_\mathrm{tot}$, is

$$E_\mathrm{tot} = 1 - e^{-\frac{3\eta_\mathrm{tot}\alpha L}{2(1-\alpha)d_\mathrm{g}}}. \qquad \text{Eq. S28}$$

Although the grain size distribution and porosity are well characterized for AF-Fe, to our knowledge there are no simple equations to calculate efficiency of a polydisperse grain size distribution. Porosity due to packing lognormally distributed diameters of solid spheres has been calculated previously and tends to decrease for wider distributions.[12] The size distribution of grains in this study is shown in **Figs. 1c** & **S9** revealing a very broad distribution over orders of magnitude following more of a power law dependence rather than a lognormal distribution. Much of the grain volume in the AF-Fe is due to $d_\mathrm{g} > 0.1$ mm, and the vast majority of grain numbers are due $d_\mathrm{g} < 0.1$ mm (**Figs. S9**). We, therefore, calculated an equivalent single grain size, $d_\mathrm{g} = 0.139 \pm 0.003$ mm, based on the measured pressure drop, $\Delta P = 0.029 \pm 0.001$ bar, that satisfies Ergun's equation[13] following

$$f_v = 1.75 + \frac{150\alpha}{Re},$$ Eq. S29

where the friction factor, $f_v$, is given by

$$f_v = \frac{\Delta P d_{\mathrm{g}}(1-\alpha)^3}{L\rho_{\mathrm{air}}U^2\alpha}.$$ Eq. S30

Using this approach to estimate filtration efficiencies has to be considered with the following caveats in mind. In our calculations, we assumed spherical monodispersed grains, while our material as previously mentioned is irregular in shape and has a very broad size distribution. Therefore, we expect non-ideal filtration behaviour and error in our calculation of filter efficiency. In addition, parameters such as surface roughness, charge, and surface hydrophilicity of the AF-Fe may contribute additional mechanisms for trapping aerosols. Therefore, while calculating filtration efficiencies using the ideal assumption of spherical particles can give valuable information, it should be treated as a qualitative result rather than a quantitative one.

#### *S1.13 Data analysis and figures*

All data analysis and plots were prepared using Matlab_R2020b on Mac OS 10.14.2. Figures and schematics were compiled using Adobe Illustrator.

**References**


1. Woo, M. H., Hsu, Y. M., Wu, C. Y., Heimbuch, B. & Wander, J. Method for contamination of filtering facepiece respirators by deposition of MS2 viral aerosols. *Journal of Aerosol Science* **41**, 944–952 (2010).
2. Kropinski, A. M., Nazzocco, A., Waddell, T. E., Lingohr, E. & Johnson, R. P. Enumeration of Bacteriophages by Double Agar Overlay Plaque Assay Andrew. in *Bacteriophages: Methods and Protocols* (eds. Clokei, M. R. J. & Kropiniski, A. M.) vol. 1 69--76 (Springer, 2009).
3. Block, K. A. *et al.* Disassembly of the cystovirus φ{symbol}6 envelope by montmorillonite clay. *Microbiologyopen* **3**, 42–51 (2014).
4. Palika, A. *et al.* An antiviral trap made of protein nanofibrils and iron oxyhydroxide nanoparticles. *Nature Nanotechnology* **16**, 918–925 (2021).
5. Brauer, R. & Chen, P. Influenza virus propagation in embryonated chicken eggs. *Journal of Visualized Experiments* **2015**, 1–6 (2015).
6. Spearman, C. The method of 'right and wrong cases (constant stimuli) without Gauss's formulae. *British journal of psycholog* **2**, 227–242 (1908).
7. Kärber, G. Beitrag zur kollektiven Behandlung pharmakologischer Reihenversuche. *Naunyn-Schmiedebergs Archiv für experimentelle pathologie und pharmakologie* **162**, 480–483 (1931).
8. Hierholzer, J. & Killington, R. Virus isolation and quantitation. in *Virology methods manual* (eds. Mahy, B. W. & Kangro, H. O.) 25–46 (Academic Press, 1996).
9. Washburn, E. The dynamics of capillary flow. *Physical Review* **17**, 273–283 (1921).
10. Otani, Y., Kanaoka, C. & Emi, H. Experimental Study of Aerosol Filtration by the Granular Bed Over a Wide Range of Reynolds Numbers. *Aerosol Science and Technology* **10**, 463–474 (2007).
11. *Aerosol Measurement: Principles, Techniques, and Applications, Third Edition*. (John Wiley & Sons, 2011). doi:10.1007/978-94-010-0786-3_5.
12. Farr, R. S. & Groot, R. D. Close packing density of polydisperse hard spheres. *The Journal of Chemical Physics* **131**, 244104 (2018).
13. Ergun, S. Fluid flow through packed columns. *Chem Eng Prog* **48**, 89–94 (1952).

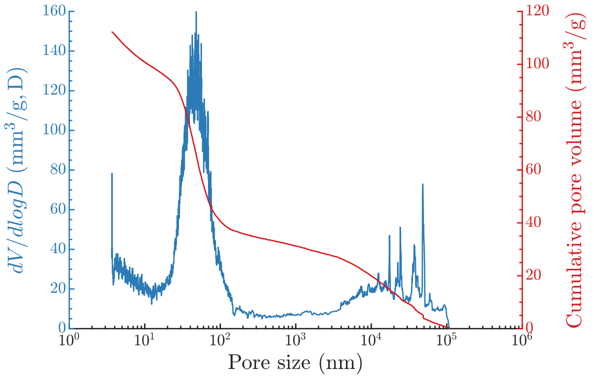


**Figure S1. Intra-particle pore size distribution of AF-Fe determined using mercury intrusion porosimetry (MIP).** The pore size distribution (left axis) and cumulative pores size distribution (right axis) show pores in size ranges of tens and thousands of nanometers.

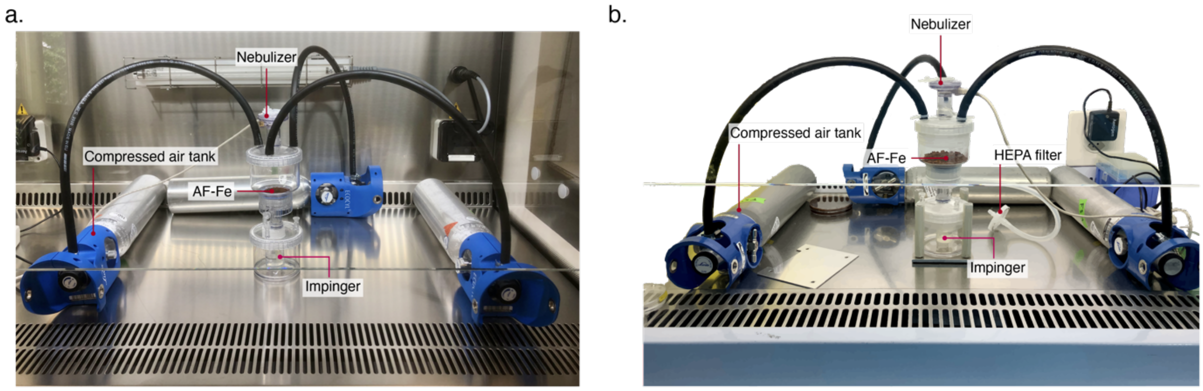


**Figure S2. Images of the experimental setup.** (a) Photo of the experimental setup in a laminar flow hood. This setup was used to run experiments with bacteriophages. (b) Vectorized image of the experimental setup in a laminar flow hood; this setup was used to run experiments with human viruses. Compared to the setup in panel (a), here it is complemented with sealings for all connections and a HEPA filter at the outlet. The back-wall of the hood in the image was deleted for clarity.

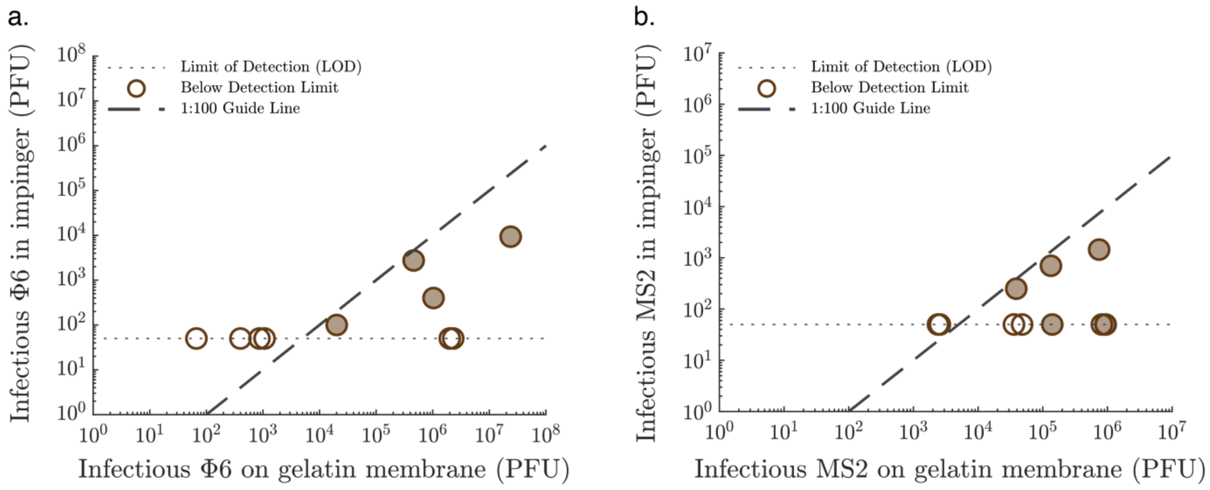


**Figure S3. The efficiency of the gelatine membranes in trapping virus-loaded aerosols.** Infectious Φ6 (a) and MS2 (b) passing through the gelatine membranes, i.e., trapped by the impinger, versus infectious viruses captured on the gelatine membrane. The capture efficiency of the gelatine membranes exceeded 99% in all cases.

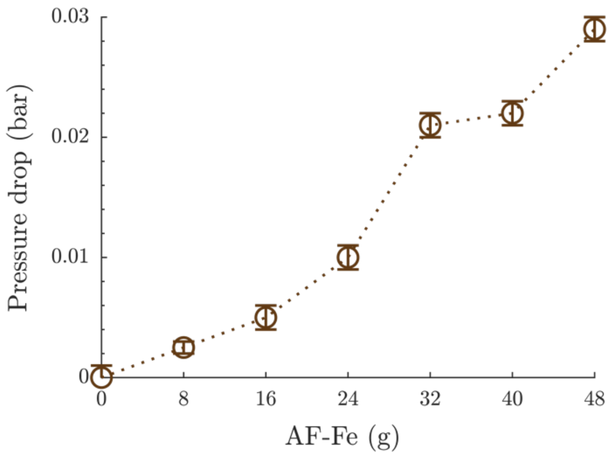


**Figure S4. Pressure drop across different amounts of AF-Fe.** The pressure drop measured versus different amounts of AF-Fe in the same setup as the one used for aerosol filtration experiments. 48 g of AF-Fe corresponds to ≈ 2 cm thickness and 48 ml volume of AF-Fe. Unless otherwise mentioned, aerosol filtration experiments were conducted using 48 g of AF-Fe.

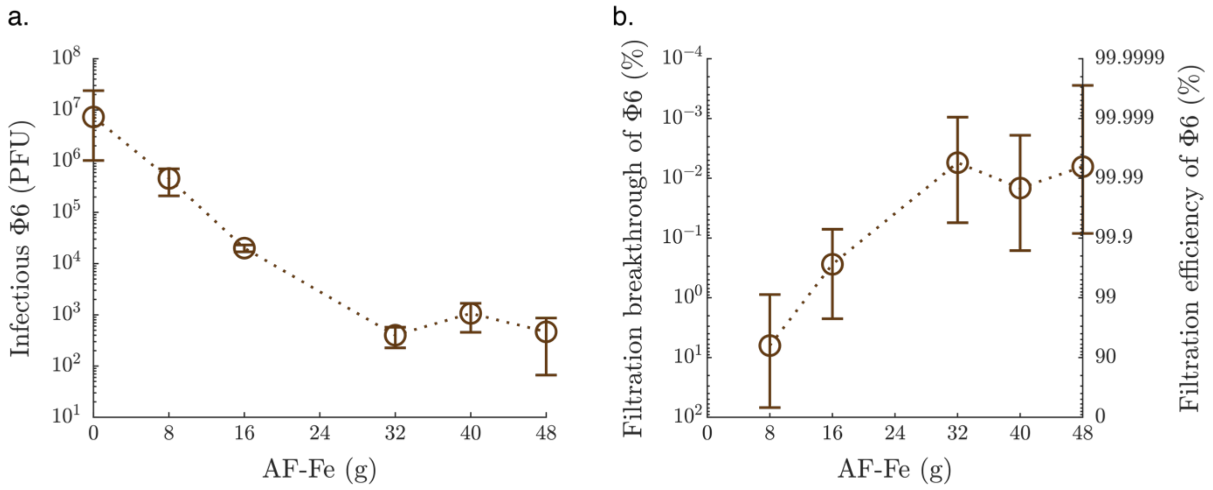


**Figure S5. Filtration efficiency of Φ6 versus the amount of AF-Fe.** (a) Infectious Φ6 recovered from the gelatine membranes versus the amount of AF-Fe used for filtration. The plotted values for 8, 16, 32, and 40 g AF-Fe represent the average from one experiment with three technical replicas; the error bars are equal to the standard deviation. For the control experiment, i.e., the experiment with 0 g AF-Fe, the plotted value represents the average from four replicas, each with three technical replicas, with the error bars representing the range, i.e., the maximum and minimum values from the four replicas. For the experiment with 48 g AF-Fe, the plotted value represents the average from two replicas, each with three technical replicas, with the error bars representing the range. (b) Filtration breakthrough and efficiency of Φ6 versus the amount of AF-Fe used. The plot values are based on the infectivity shown in panel (a). The error bars represent the maximum and minimum efficiencies by considering the extreme values in panel (a), i.e., dividing the minimum infectivity of each experiment by the maximum of the control (at 0 g AF-Fe) and vice versa.

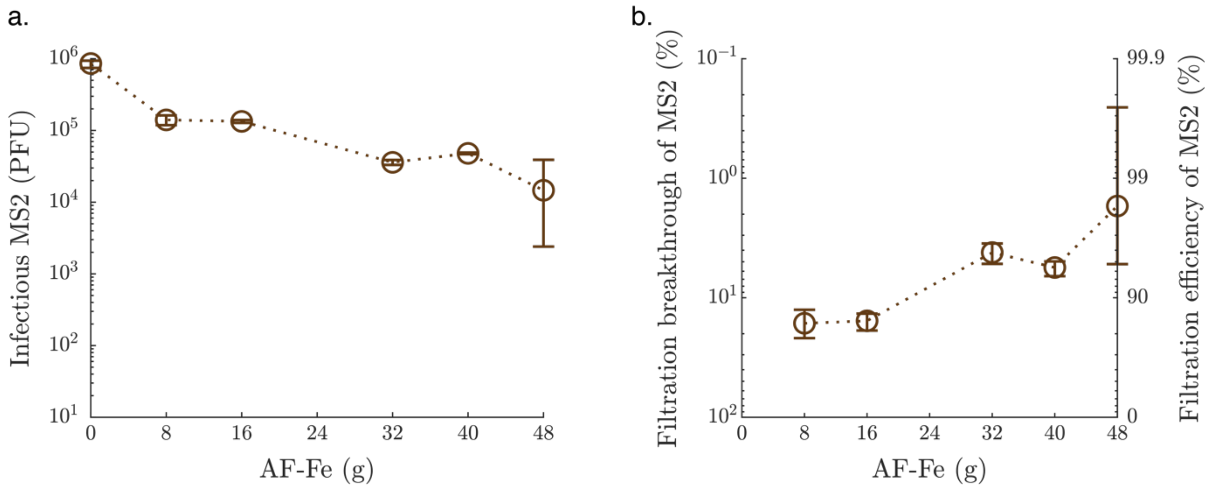


**Figure S6. Filtration efficiency of MS2 versus the amount of AF-Fe.** (a) Infectious MS2 recovered from the gelatine membranes versus the amount of AF-Fe used for filtration. The plotted values for 8, 16, 32, and 40 g AF-Fe represent the average from one experiment with three technical replicas; the error bars are equal to the standard deviation. For the control experiment, i.e., the experiment with 0 g AF-Fe, the plotted value represents the average from four replicas, each with three technical replicas, with the error bars representing the range, i.e., the maximum and minimum values from the four replicas. For the experiment with 48 g AF-Fe, the plotted value represents the average from three replicas, each with three technical replicas, with the error bars representing the range. (b) Filtration breakthrough and efficiency of MS2 versus the amount of AF-Fe used. The plot values are based on the infectivity shown in panel (a). The error bars represent the maximum and minimum efficiencies by considering the extreme values in panel (a), i.e., dividing the minimum infectivity of each experiment by the maximum of the control (at 0 g AF-Fe) and vice versa.

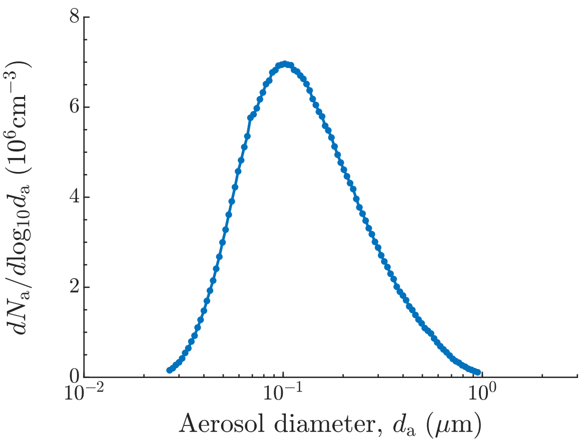


**Figure S7. Aerosol size distribution.** Size distribution, $dN_a/d\log_{10}d_a$, for aerosolized artificial saliva/mucin solution produced using the Aeroneb® Lab nebulizer unit (Kent Scientific, U.S.A.) with nominal aerosol size range 4 – 6 µm. The artificial saliva/mucin solution was used for all Φ6 and MS2 filtration experiments. $d_a$ is the aerosol diameter, $d\log_{10}d_a$ is the width of a $d_a$ bin on a logarithmic scale and $dN_a$ is the number of particles in a size bin per cm$^3$ of air. Although the nebulizer generates particles with $d_a$ between 4 and 6 µm, they quickly evaporate in air stream resulting in an equilibrium relative humidity, *RH*, between 80% and 85%. This explains why aerosols with $d_a < 1$ µm are abundant.

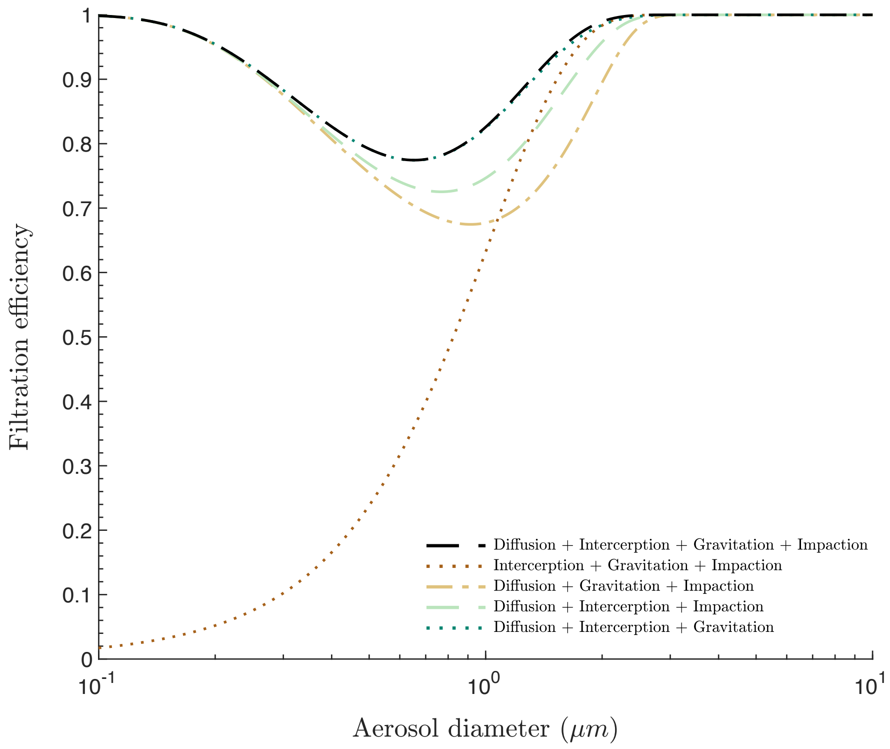


**Figure S8. Total filtration efficiencies from different contributions.** Total filtration efficiencies across a grain bed filter with monodispersed grain size $d_g = 0.139$ mm. The efficiencies were calculated considering different combinations of contributions from diffusion, interception, gravitational settling, and impaction.

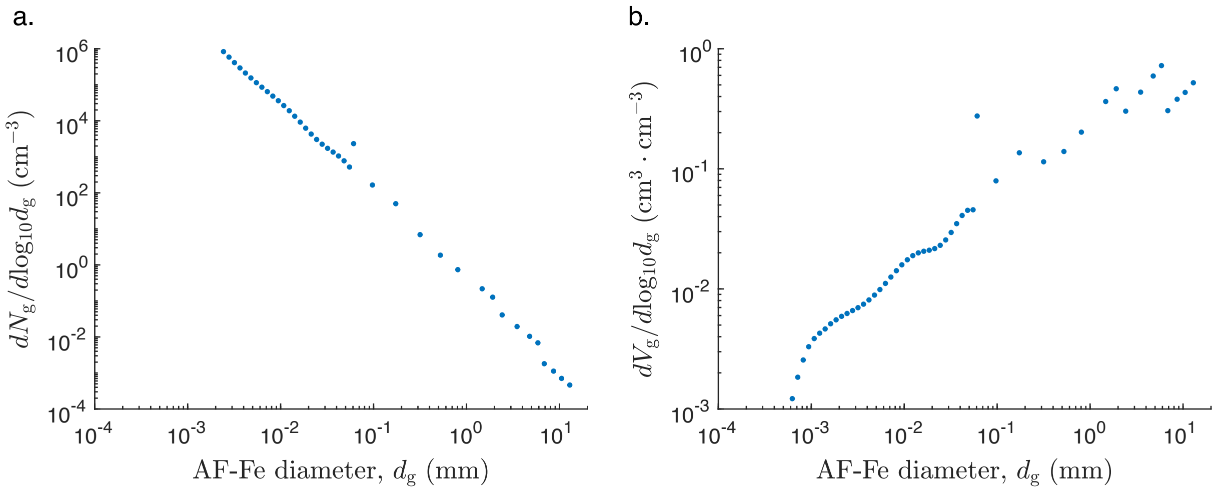


**Figure S9. Number and volume size distribution of AF-Fe.** (a) Number, $dN_g/d\log_{10}d_g$, and (b) volume, $dV_g/d\log_{10}d_g$, distributions for grains in the granular bed filter. $d_g$ is the grain diameter, $d\log_{10}d_g$, is the width of a $d_g$ bin on a logarithmic scale, and $dN_g$ and $dV_g$ are the number and volume, respectively, of particles in a size bin per cm$^3$ of the filter.
